# Extracellular Vesicles Derived from *L-MYC* Neural Stem Cells Mediate Neuroprotection in 3D Models of Chemotherapy- and Radiation-Induced Neurotoxicity

**DOI:** 10.64898/2026.08.26.747380

**Authors:** Lance G. A. Nunes, Isabella Vasquez, Brecken Enright, Liv Chen, Sunita K. Patel, Russell C. Rockne, Stephanie Yoon, Margarita Gutova

**Affiliations:** Department of Stem Cell Biology and Regenerative Medicine, Beckman Research Institute, City of Hope, Duarte, CA 91010, USA; (L.G.A.N.); (I.V.); (B.E.); (L.C.); Department of Population Sciences and Supportive Care Medicine, Beckman Research Institute, City of Hope, Duarte, CA 91010, USA; Department of Computational and Quantitative Medicine, Division of Mathematical Oncology, Beckman Research Institute, City of Hope, Duarte, CA 91010, USA; Department of Radiation Oncology, Beckman Research Institute, City of Hope, Duarte, CA 91010, USA

**Keywords:** cancer therapy-related cognitive impairment (CRCI), chemotherapy-induced neurotoxicity, radiation-induced brain injury, L-MYC-immortalized neural stem cells (LMNSCs), LMNSC-derived extracellular vesicles (LMNSC-EVs), neuroprotection, neurorestoration, regenerative medicine, 3D human neural tissue models

## Abstract

**Background/Objectives:** Cancer survivors frequently experience long-term neurocognitive impairments following chemotherapy and cranial irradiation, yet experimental models that enable mechanistic investigation of therapy-induced neurotoxicity at the transcriptional level remain limited. This study aimed to develop a human three-dimensional (3D) neural tissue model derived from L-MYC-immortalized neural stem cells (LMNSCs) and use transcriptomic profiling to identify molecular pathways underlying chemotherapy- and radiation-induced neural injury and extracellular vesicle (EV)-mediated recovery.

**Methods:** LMNSCs were differentiated in a 3D, methylcellulose-based culture to generate neural tissue containing neurons, astrocytes, and oligodendrocytes in the absence of microglia. Cultures were exposed to methotrexate (MTX) or ionizing radiation to induce neural injury and subsequently treated with LMNSC-derived EVs. Neural injury and repair mechanisms were evaluated by immunocytochemistry and bulk transcriptomics.

**Results:** MTX and irradiation induced dose-dependent injury, exhibited by loss of neuronal complexity and reduced glial populations. LMNSC-EV treatment promoted recovery of neuronal and glial populations following MTX- and irradiation-induced injury. Transcriptomic analysis of irradiated cultures revealed activation of inflammation, DNA damage, and stress-response pathways, which were attenuated after treatment with LMNSC-EVs.

**Conclusions:** LMNSC-based 3D neural tissue provides a platform for modeling cancer therapy-induced neurotoxicity. Furthermore, LMNSC-EVs represent a promising cell-free regenerative therapeutic that modulates injury-associated inflammatory, stress, and metabolic transcriptional programs after radiation-induced neural injury.

**Simple Summary:** Many cancer survivors experience persistent problems with memory, attention, and learning after chemotherapy or radiotherapy, yet the biological mechanisms underlying these cognitive side effects remain poorly understood. Progress has been limited by the lack of human laboratory models that accurately reproduce treatment-induced brain injury. To address this need, we developed a three-dimensional human neural tissue model containing immature neurons, astrocytes, and oligodendrocytes derived from L-MYC-immortalized human neural stem cells, lacking microglia. This platform was used to model cancer therapy-induced neural injury and evaluate the regenerative potential of neural stem cell-derived extracellular vesicles (EVs)—small bioactive particles released by stem cells. We found that cancer therapies caused significant injury to neuronal, oligodendroglial, and astrocytic cell populations, whereas EV treatment promoted recovery of these cells and activated biological pathways associated with tissue repair. These findings establish a reproducible human neural tissue platform for studying cancer therapy-related neurotoxicity and support neural stem cell-derived EVs as a promising regenerative therapeutic strategy to prevent or reduce treatment-induced cognitive impairment.

## 1. Introduction

Advances in cancer therapy have significantly improved survival across many malignancies; however, long-term neurological complications remain a major clinical challenge [1]. A growing number of cancer survivors experience persistent cognitive deficits collectively referred to as cancer therapy-related cognitive impairment (CRCI)[2,3]. These deficits may affect memory, attention, executive function, and processing speed and can persist for months or years after treatment, significantly impacting quality of life and functional outcomes in both adult and pediatric cancer survivors [4–7]. Both chemotherapy and cranial irradiation contribute to CRCI through mechanisms that disrupt multiple cellular components of the central nervous system. The chemotherapeutic agent methotrexate (MTX) is widely used in the treatment of hematologic malignancies in children. Although effective in treating cancer, MTX often induces neurotoxicity through oxidative stress, mitochondrial dysfunction, neuroinflammation, and impaired neurogenesis [8,9]. MTX exposure has been associated with damage to neural progenitor cells and oligodendrocyte precursor cells (OPCs), resulting in alterations in white matter integrity and neuronal signaling [8,10]. Similarly, cranial radiotherapy remains a critical and effective component of treatment for many primary and metastatic brain tumors, yet radiation exposure can damage neural stem cell (NSC) niches, vascular structures, and glial populations, leading to chronic neuroinflammation, progressive neurodegeneration, and long-term cognitive decline [11–13].

Despite the increasing recognition of CRCI, the biological mechanisms underlying cancer therapy-induced neurotoxicity remain incompletely understood. A major limitation that remains is the lack of experimental models that accurately model human neural compartment injury and repair. Traditional two-dimensional neural cultures do not capture multicellular interactions and tissue architecture, while animal models often fail to fully reproduce human-specific responses to cancer therapy. Consequently, there remains a critical need for reproducible, human neural tissue models for mechanistic studies and therapeutic development that improve the predictive value of preclinical studies [14]. Recent advances in three-dimensional (3D) neural culture systems derived from human stem cells provide new opportunities to address these limitations [15,16]. Human NSC-derived spheroids and organoid models, including those derived from L-MYC-immortalized human NSCs (LMNSCs), generate multicellular neural tissues containing neuronal, astrocytic, and oligodendroglial lineage cells within a structured microenvironment [17–19]. These 3D multicellular systems provide a valuable platform for investigating mechanisms underlying neural compartment injury, intercellular interactions, and responses to regenerative therapies in a more structurally relevant context compared to conventional approaches.

In parallel with advances in disease modeling, regenerative approaches aimed at restoring neural function after injury have attracted considerable attention. NSCs have demonstrated the capacity to promote brain repair through multiple mechanisms, including modulation of inflammation, secretion of neurotrophic factors, and stimulation of endogenous regenerative pathways [20–23]. Recent studies using intranasally delivered LMNSCs demonstrated migration to sites of brain injury, improved cognitive functional recovery, and modulation of inflammatory pathways, supporting their therapeutic potential in brain injury [23,24]. Increasing evidence suggests that many of the therapeutic effects of stem cells are mediated by paracrine signaling, particularly through extracellular vesicles (EVs) [21,25–30]. EVs carry diverse bioactive cargo, including proteins, lipids, mRNAs, and microRNAs, which can influence cellular signaling pathways involved in neuronal survival, synaptic plasticity, and tissue repair. NSC-derived EVs have shown promising neuroprotective effects in preclinical models of neurological injury, including traumatic brain injury and neurodegenerative disorders, highlighting their potential as a cell-free therapeutic approach [29,31]. Despite progress in both disease modeling and EV-based therapeutics, relatively few studies have combined these approaches to investigate chemotherapy- and radiation-induced brain injury. Developing such systems is critical for improving our understanding of cancer therapy-induced neurotoxicity and for identifying interventions that can mitigate neurological damage in cancer patients.

To address this need, we developed a human 3D neural tissue model derived from LMNSCs that generate neuronal, astrocytic, and oligodendroglial populations, resembling those found in the human brain. Using this 3D platform, we modeled neurotoxicity induced by MTX and ionizing radiation and investigated whether LMNSC-derived EVs (LMNSC-EVs) could promote recovery of neural populations and restore protective molecular pathways. By integrating a human 3D neural tissue culture system with regenerative EV-based therapy, this study establishes a translational platform for investigating mechanisms of therapy-induced brain injury and for evaluating potential neuroprotective strategies relevant to cancer survivors. We hypothesized that a human 3D neural tissue model derived from LMNSCs can recapitulate key features of chemotherapy- and radiation-induced neurotoxicity and serve as a platform to evaluate regenerative interventions. Specifically, we sought to determine whether exposure of 3D neural cultures to MTX and ionizing radiation induces lineage-specific neural injury and transcriptional alterations. In addition, we evaluated whether EVs of known composition derived from LMNSCs [25] could mitigate therapy-induced damage and promote recovery of neural cell populations and neurorestorative signaling pathways. Together, these studies establish a human-relevant experimental system for modeling cancer therapy-induced neural injury and for identifying potential neuroprotective and neurorestorative strategies for cancer survivors.

## 2. Materials and Methods

### 2.1. Expansion of LMNSC01 and LMNSC02 Cells

LMNSC01 and LMNSC02 cells were expanded and propagated using a previously described protocol [19,25]. To expand LMNSCs, cells were cultured in T-175 flasks using NeuroCult NS-A Basal Medium (Human) (StemCell Technologies, Cat# 05750) and supplemented with NeuroCult NS-A Proliferation Supplement (Human) (Cat# 05751). The cells were further supplemented with heparin (0.2 × 10^−^⁴% w/v; Cat# 07980), basic fibroblast growth factor (bFGF, 10 ng/mL; Cat# 78003), and epidermal growth factor (EGF, 10 ng/mL; Cat# 78006). Cells were maintained under physiological hypoxia (4% O_2_, 5% CO_2_) at 37 °C to optimize growth. Isolation and propagation procedures were conducted in accordance with approved protocols at City of Hope.

### 2.2. Generation of 3D LMNSC01 and LMNSC02 Cultures

3D cultures were generated using (50%) StemXVivo Methylcellulose Concentrate (MC) (R&D Systems, Cat#HSC011) and (50%) NeuroCult NS-A Basal Medium (Human) (NC) (StemCell Technologies, Cat# 05750) supplemented with either NeuroCult NS-A Differentiation Supplement (Human) (Cat# 05752) or NeuroCult NS-A Proliferation supplement (Human) with heparin, EGF, and bFGF. Briefly, 5 × 10^4^ cells/cm^2^ were suspended in 250 µL of NeuroCult Differentiation medium (NC-diff) or 2.5 × 10^4^ cells/cm^2^ were suspended in 250 µL of NeuroCult Proliferation medium (NC-prolif) and were carefully mixed into another 250 µL of MC (MC:NC-prolif (1:1)). This 500 µL suspension was then added to a 24-well plate using an 18-gauge needle and 1 mL syringe. The cells were maintained under physiological hypoxia (4% O_2_) at 37 °C throughout the experiment, where 50 µL of fresh medium was added every other day to the top of the wells.

### 2.3. Irradiation Exposure of 3D Cultures and Treatment with LMNSC-EVS

For cells in MC:NC (1:1) proliferation cultures, cells were irradiated on day 5. For cells in MC:NC-diff (1:1) cultures, cells were irradiated on day 8. Radiation doses of 0, 4, 8, and 12 Gy were selected based on preliminary dose–response studies performed in our laboratory and pilot transcriptomic analyses of irradiated LMNSC-derived organoids (Figure S2). All cells were irradiated at 0 Gy, 4 Gy, 8 Gy, and 12 Gy using the MultiRad 160 (Precision X-Ray Irradiation, Madison, US). Cultures were irradiated using the following settings: (Dose 1 Gy/min, FSD 37 cm, Filter: Copper 0.30 mm, kV 160, mA 13.3). Immediately after irradiation, fresh medium containing LMNSC-EVs (2 µg/mL) was added to the top of the wells. The 3D cultures were then left for an additional 4 days to recover, during which 50 µL of fresh medium was added every other day to the top of the wells. Cells in MC:NC-proliferation (1:1) cultures were cultured for a total of 9 days, while cells in MC:NC (1:1) differentiation cultures were cultured for a total of 12 days.

### 2.4. MTX Exposure of 3D Cultures and Treatment with LMNSC-EVS

For cells in MC:NC (1:1) proliferation cultures, cells were treated with MTX on day 4. For cells in MC:NC-diff (1:1) cultures, cells were treated with MTX on day 7. MTX concentrations (0–2 µM) were selected based on published in vitro studies of MTX-induced neural injury and glial activation [8], prior work from our laboratory using LMNSC-derived neural organoids, and preliminary dose-finding studies performed in the present culture system. MTX was supplied as a liquid and diluted directly into the culture medium to achieve the indicated final concentrations. All cultures were provided fresh medium containing MTX at the final concentrations of 0 µM, 1 µM, and 2 µM, and were left to incubate for 3 days. After this incubation period, cells were then provided with fresh medium containing LMNSC-EVs (2 µg/mL). Cells were then left to recover for an additional 4 days, during which 50 µL of fresh medium was added every other day to the top of the wells. Cells in MC:NC-prolif (1:1) cultures were cultured for a total of 11 days, while cells in MC:NC-diff (1:1) cultures were cultured for a total of 14 days.

### 2.5. Isolation of LMNSC-EVs

LMNSC01- and LMNSC02-EVs were isolated using a previously described protocol [25]. Briefly, LMNSCs were cultured in T-175 flasks at a high cell density (10^7^ cells) for 3 days in NC-prolif containing heparin, EGF, and bFGF. The conditioned medium (CM) was collected and centrifuged at 500g for 10 min to pellet debris. Afterwards, CM was concentrated to 500 µL using a Vivaspin 100 kDa MWCO concentrator (Satorius, Cat# PNVS2042) [25]. LMNSC-EVs were then isolated in PBS using an Izon-qEV original column, 70 nm Gen 2 (IZON, Cat# ICO-70-12745), and further concentrated with a Vivaspin 100 kDa MWCO concentrator to create a working volume. Protein concentration was measured using Pierce BCA Protein Assay Kit (Thermo Scientific, Waltham, MA, USA, Cat# 23227) according to the manufacturer’s recommendation.

### 2.6. Generation of 2D LMNSC Cultures

For bulk RNA sequencing, 2D cultures were generated. LMNSCs were plated at a concentration of 5 × 10^4^ cells/cm^2^ in each well of a 6-well plate, in 2 mL of NC-prolif supplemented with heparin, EGF and bFGF. LMNSCs were allowed to attach for 24 hours, and the resulting medium was replaced with 2 mL of NC-diff. Half medium changes were performed every other day, and the medium was completely replaced every week, before cultures were exposed to MTX or irradiation. For cells exposed to irradiation, a complete medium change was performed right after irradiation, where select cultures were treated with LMNSC-EVs (2 µg/mL). For cells exposed to MTX, a complete medium change was performed 3 days after exposure, where select cultures were treated with LMNSC-EVs (2 µg/mL). Both NC-diff cultures treated with LMNSC-EVs were left to incubate for an additional 4 days before cells were collected for RNA isolation.

### 2.7. RNA Isolation of 2D LMNSC Cultures

To evaluate RNA concentration and purity, RNA was extracted using the RNeasy Mini kit (Qiagen, Cat# 74104) in accordance with the manufacturer’s protocol, with an on-column DNase digestion step using the RNase-Free DNase Set (Qiagen, Cat#79254). Due to excess phenol and guanidine contamination after RNA clean-up, an ethanol precipitation step was also performed using a previously described protocol [32]. Briefly, sodium acetate (3M, 5.2 PH) was diluted to 0.3M in isolated RNA suspended in nuclease-free water. 3 volumes of 100% ice-cold ethanol were then added to the RNA samples and left to precipitate overnight at 4 °C. Supernatant was removed using centrifugation at 4 °C, and precipitated RNA was washed twice with 70% ice-cold ethanol. The resulting ethanol was removed, and precipitated RNA pellets were suspended in nuclease-free water. RNA concentration and purity were examined using NanoDrop OneC (Thermo Scientific).

### 2.8. RNA-Seq Analysis of 2D LMNSC Cultures

Total RNA isolated from NC-diff cultures was sequenced at Plasmidsaurus (Louisville, KY). Raw base call (BCL) files were converted to FASTQ format and demultiplexed using BCL Convert (v4.3.6) and fqtk (v0.3.1). Reads were processed with FastP (v0.24.0) for poly-X trimming, 3′ quality trimming, enforcement of a minimum Phred quality score of 15, and a minimum read length of 50 bp. Filtered reads were aligned to the GRCh38 reference genome using the STAR aligner (v2.7.11) with non-canonical splice junction removal and unmapped read output enabled. Resulting BAM files were coordinate-sorted using samtools (v1.22.1), and PCR and optical duplicates were removed using UMICollapse (v1.1.0) based on unique molecular identifiers (UMIs). Alignment quality metrics, strand specificity, and genomic feature distribution were assessed using RSeQC (v5.0.4) and Qualimap (v2.3), and a comprehensive quality control report was generated with MultiQC (v1.32). Gene-level quantification was performed using featureCounts (Subread package v2.1.1) with strand-specific counting, fractional assignment of multi-mapping reads, and annotation of exons and 3′ UTR regions grouped by gene_ID. Final count matrices were annotated with gene biotype and metadata extracted from the reference GTF file. Differential gene expression analysis was conducted using edgeR (v4.0.16), with lowly expressed genes filtered using edgeR::filterByExpr under default parameters. Functional enrichment was performed using gene expression analysis with gseapy v0.12 using the MSig DB Hallmark gene set.

### 2.9. Immunocytochemistry of 3D Brain Cultures

Immunocytochemistry (ICC) was performed after the endpoint of irradiation and MTX experiments. MC was carefully diluted with additional NC-diff medium and removed from culture. Afterwards, cells were fixed with 4% PFA for 30 min. After fixation, cells were washed three times in ten-minute intervals with 1× TBS including 0.1% Triton X-100. After washes, cells were blocked with CAS Block (ThermoFisher, Cat# 008120) for one hour. Primary antibodies were incubated with cells overnight at 4 °C. Primary antibodies used included β-tubulin III (BioLegend, Cat# 801202; 1:500), GFAP (Abcam, Cat# 53554; 1:500) and OLIG1 (Invitrogen, Cat# PA5-21613; 1:150) to analyze neuronal, astrocytic, and oligodendrocyte markers retrospectively. After incubation, three additional washes were performed with 1× TBS with 0.1% Triton to minimize non-specific staining. Secondary antibodies were incubated for two hours at room temperature. Secondary antibodies used include Alexa Fluor 488 donkey anti-goat (Invitrogen, A21467; 1:1000), Alexa Fluor 555 donkey anti-rabbit (Invitrogen, A31572; 1:1000), and Alexa Fluor 647 donkey anti-mouse (Invitrogen, A32787; 1:1000). Hoechst (Invitrogen, H3570; 1:1000) was used as a counterstain to visualize nuclei and identify cells. Three additional washes were performed with 1× TBS with 0.1% Triton to minimize background signal. Z-stack imaging was performed using Zeiss Observer II (Zeiss, Göttingen, Germany) at 10×. A total of 5 z-stacks were taken for each image, with a 5 µm z-step size. Maximum projection and background subtraction were performed to minimize noise using Zeiss ZEN (ZEN, VER 3.12).

### 2.10. Analysis of OLIG1 Positive Cells

The proportion of OLIG1-positive cells was quantified using IMARIS Microscopy Image Analysis Software (Oxford Instruments, VER 11.0.1). Data from two independent experiments were combined for analysis, where the first experiment contained two biological replicates with one technical replicate each, while the second experiment contained three biological replicates with three technical replicates each. Background subtraction was performed in Zeiss ZEN, and files were then converted to native IMARIS files. First, Hoechst-positive cells were converted into surfaces to identify nuclei, while cells stained for OLIG1 were converted into spots to identify OLIG1-positive cells using the algorithm for the quality metric. Double positives and false positives were manually removed, and raw data were exported. Total cell count was analyzed using two-way ANOVA, while the percentage of OLIG1-positive cells was analyzed using one-way ANOVA.

### 2.11. Sholl Analysis

Neuronal complexity was quantified using Sholl analysis in Fiji (ImageJ). Data from two independent experiments were combined for analysis, where the first experiment contained one biological replicate with one technical replicate, while the second experiment contained three biological replicates with three technical replicates each. Z-stack images were acquired using a Zeiss Observer II microscope (CZI format) (Zeiss, Oberkochen, Germany). CZI files were imported into FIJI (ImageJ version 1.54p), and z-stacks were projected into two-dimensional images using an average-intensity projection. Individual fluorescence channels were separated, and the β-Tubulin III channel was selected for analysis. Images were converted to binary masks using a lower threshold value of 220. Sholl analysis was then performed using the Interactive Sholl Analysis function in the FIJI Neuroanatomy plugin. The center of each neuronal cluster was manually defined using a region of interest (ROI), and concentric circles were generated beginning at a radius of 10

µm with a 10 µm step size. The maximum analysis radius was adjusted for each cluster to encompass all neuronal processes while excluding branches originating from neighboring clusters. The number of process intersections at each radius was recorded.

## 3. Results

### 3.1. Generation of LMNSC01 and LMNSC02 3D NSC-Derived Brain Tissues

To establish a human 3D neural tissue platform, LMNSCs (LMNSC01 and LMNSC02) were cultured in a 3D MC matrix mixed at a 1:1 ratio with NeuroCult NS-A Basal Medium supplemented with either NeuroCult-differentiation (NC-diff) or proliferation (NC-prolif) supplement and assessed by fluorescent ICC (Figure 1). When maintained for 14 days in MC:NC-diff (1:1) medium, both LMNSC lines formed dense 3D cellular aggregates containing multiple neural lineages (neuronal, astrocytic, and oligodendroglial). While LMNSC01 aggregates were comparatively compact and exhibited moderate branching, LMNSC02 aggregates were highly arborized, exhibiting a higher degree of neural complexity. Immunostaining revealed robust expression of neuronal biomarker β-Tubulin III, indicating the presence of differentiating neurons, alongside GFAP+ astrocytes and OLIG1+ OPCs. Hoechst nuclear staining confirmed widespread cellular distribution throughout the 3D matrix.

**Figure 1.**
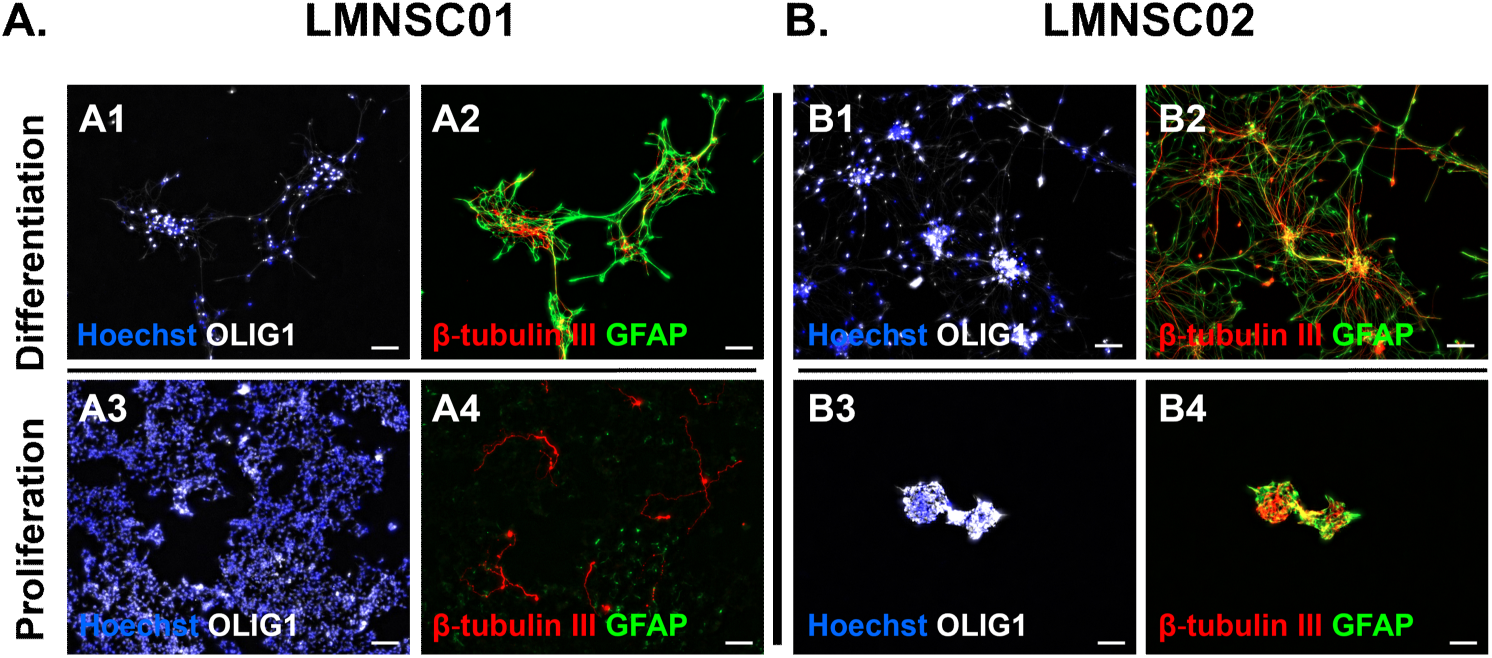
Proliferation and differentiation of LMNSC01 and LMNSC02 in 3D methylcellulose culture. (**A**) Representative images of immunocytochemistry (ICC) of LMNSC01 in an MC:NC-diff (1:1) culture for 14 days (**A1, A2**) and in MC:NC-prolif (1:1) culture for 11 days (**A3, A4**). Cells cultured in a MC:NC-diff (1:1) culture were seeded at a concentration of 5.0 × 10^4^ cells/cm^2^, while cells cultured in an MC:NC-prolif (1:1) culture were seeded at 2.5 × 10^4^ cells/ cm^2^. (**B**) Representative ICC of LMNSC02 in an MC:NC-diff (1:1) culture for 14 days (**B1, B2**) and in MC:NC-prolif (1:1) culture for 11 days (**B3, B4**). Cells cultured in an MC:NC-diff (1:1) culture were seeded at a concentration of 5.0 × 10^4^ cells/ cm^2^, while cells cultured in a MC:NC-prolif (1:1) culture were seeded at 2.5 × 10^4^ cells/cm^2^. ICC staining was done with β-Tubulin III (red), OLIG1 (white), and GFAP (green), and nuclei were stained with Hoechst dye (blue). Images were obtained on a Zeiss Observer II (10x). All images are representative maximum intensity projections from z-stack fluorescent images. Maximum projection and background subtraction were performed using Zeiss ZEN (ZEN, VER 3.12). Scale bar = 100 µm.

In parallel, LMNSCs cultured for 11 days in MC:NC-prolif (1:1) medium maintained a non-differentiated NSC phenotype, characterized by low expression of neural and glial biomarkers. Comparative analysis of LMNSC01 and LMNSC02 demonstrated consistent lineage marker expression across two donor-derived LMNSC lines, supporting the reproducibility of the 3D culture system. Together, these data demonstrate that LMNSCs proliferate and undergo multilineage neural differentiation within a 3D MC culture, generating neuronal, astrocytic, and oligodendroglial cells that model key cellular components of human neural tissue.

### 3.2. Dose-Dependent MTX-Induced Neural Injury in Differentiated 3D LMNSC Culture

To model chemotherapy-induced neurotoxicity in human 3D neural tissue, differentiated LMNSC01 and LMNSC02 cultures were exposed to increasing concentrations of MTX (0, 1, and 2 µM) and analyzed by fluorescent ICC after 14 days in MC:NC-diff (1:1) medium (Figure 2A). Untreated cultures from both LMNSC01 and LMNSC02 lines (Figure 2(B1, B2, C1, C2)) exhibited well-organized, complex multicellular organization characterized by extensive βTubulin III+ neuronal processes, GFAP+ astrocytes, and OLIG1+ OPCs.

**Figure 2.**
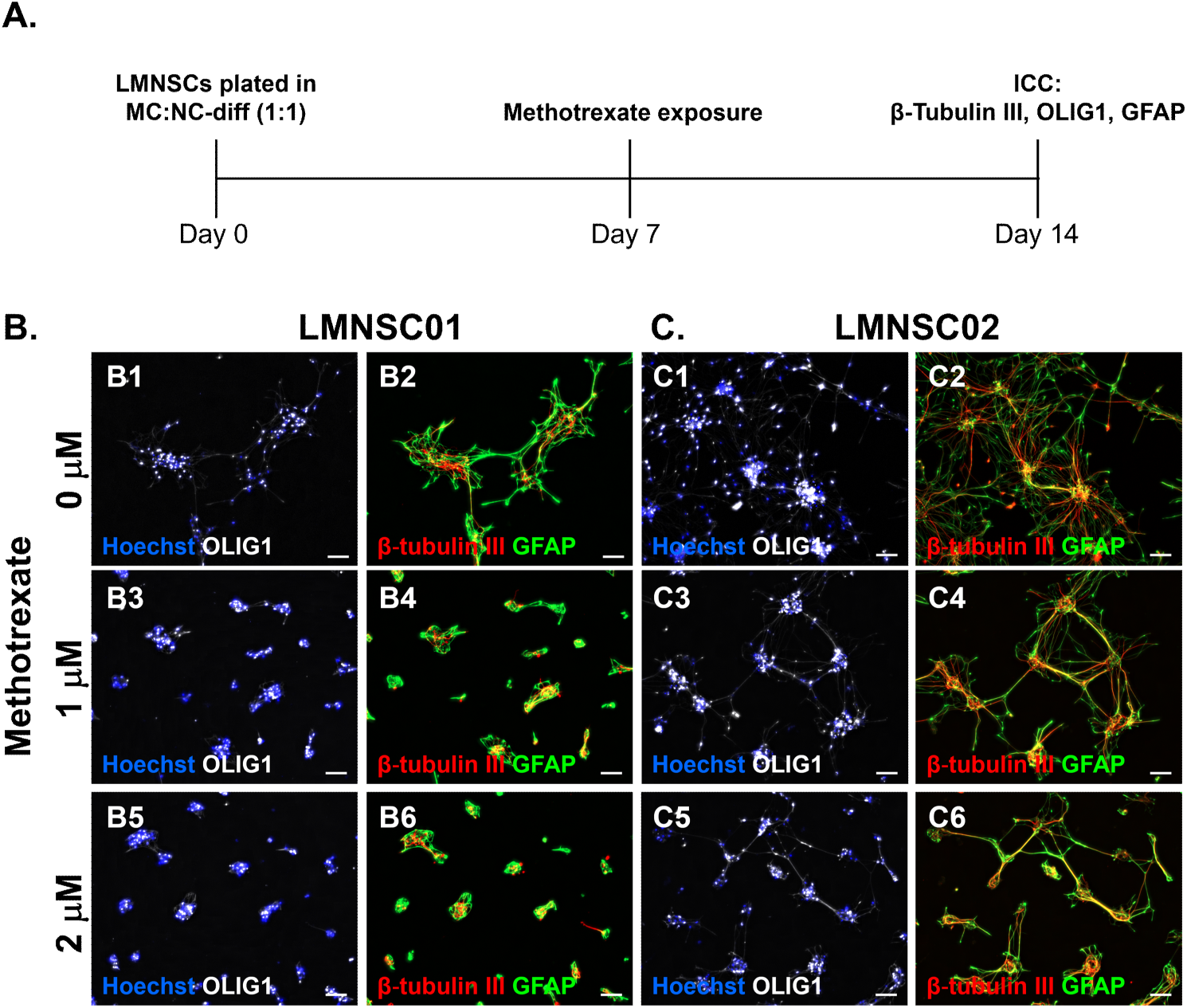
Dose-dependent MTX-induced injury in differentiated 3D LMNSC01 and LMNSC02 cultures. (**A**) Experimental timeline; (**B**) representative images of immunocytochemistry (ICC) of LMNSC01 cells exposed to methotrexate (MTX) at concentrations of 0 µM, 1 µM, 2 µM in a MC:NC (1:1) differentiation culture for 14 days; (**C**) representative images of ICC of LMNSC02 cells exposed to MTX at concentrations of 0 µM, 1 µM, 2 µM in MC:NC (1:1) differentiation culture for 14 days. Quantitative analyses of treatment-induced changes at selected MTX doses are presented in Figure 4 and Figure 5. ICC staining was done with (**B2, B4, B6, C2, C4, C6**) β-Tubulin III (red) and GFAP (green), and (**B1, B3, B5, C1, C3, C5**) OLIG1 (white), with nuclei stained with Hoechst dye (blue). Images were obtained on Zeiss Observer II (10x). All images are representative maximum intensity projections from z-stack fluorescent images. Maximum projection and background subtraction were performed using Zeiss ZEN (ZEN, VER 3.12). Scale bar = 100 µm.

MTX exposure induced a dose-dependent disruption of 3D neural architecture in both LMNSC01 and LMNSC02 cultures. At 1 µM MTX (Figure 2(B3, B4, C3, C4)), cultures displayed reduced neurite complexity, fragmentation of βTubulin III+ neuronal processes, and decreased OLIG1+ OPC populations, accompanied by altered spatial organization of GFAP+ astrocytes. These effects were more pronounced at 2 µM MTX (Figure 2(B5, B6, C5, C6)), where marked loss of neuronal networks, sparse and truncated neurites, and diminished OPC staining were observed, together with increased nuclear condensation detected by Hoechst labeling. Comparable structural and lineage-specific vulnerabilities were observed across both LMNSC lines, demonstrating reproducible MTX-induced neural injury within the 3D culture system.

### 3.3. Dose-Dependent Radiation-Induced Neural Injury in Differentiated 3D LMNSC Culture

To model radiation-induced neurotoxicity, differentiated LMNSC01 and LMNSC02 3D neural cultures were exposed to varying doses of ionizing radiation (0, 4, 8, and 12 Gy) and analyzed by fluorescent ICC after 12 days in MC:NC-diff (1:1) medium (Figure 3A). Control cultures (0 Gy; Figure 3(B1, B2, C1, C2)) from both LMNSC lines exhibited organized neural architecture with βTubulin III+ neurons, GFAP+ astrocytes, and OLIG1+ OPCs. Neuronal and astrocytic processes were highly arborized, indicative of neural differentiation throughout the 3D matrix.

**Figure 3.**
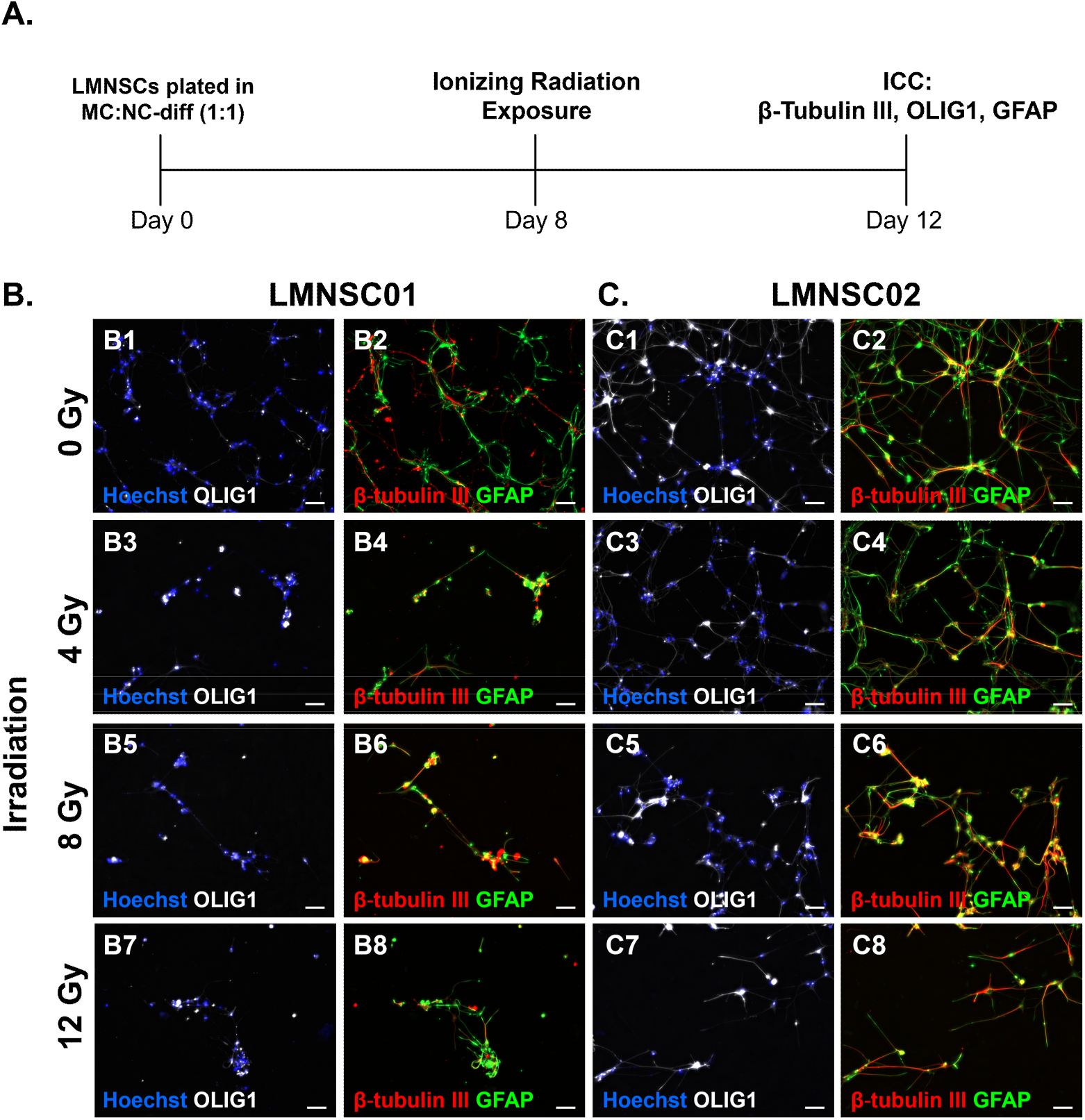
Dose-dependent radiation-induced injury in differentiated 3D LMNSC01 and LMNSC02 cultures. (**A**) Experimental timeline; (**B**) representative images of immunocytochemistry (ICC) of LMNSC01 cells exposed to 0 Gy, 4 Gy, 8 Gy, and 12 Gy ionizing radiation in a 3D MC:NC (1:1) differentiation culture for 12 days; and (**C**) representative images of ICC of LMNSC02 cells exposed to 0 Gy, 4 Gy, 8 Gy, and 12 Gy ionizing radiation in a 3D MC:NC (1:1) differentiation culture for 12 days. Quantitative analyses of radiation-induced changes in selected doses are presented in Figure 6 and Figure 7. ICC staining was done with (**B2, B4, B6, B8, C2, C4, C6, C8**) β-Tubulin III (red) and GFAP (green), and (**B1, B3, B5, B7, C1, C3, C5, C7**) OLIG1 (white) with nuclei stained with Hoechst dye (blue). Images were obtained using Zeiss Observer II (10x). All images are representative maximum intensity projections from z-stack fluorescent images. Maximum projection and background subtraction were performed using Zeiss ZEN (ZEN, VER 3.12). Scale bar = 100 µm.

Increasing doses of radiation induced progressive, dose-dependent disruption of neural structure and lineage integrity. Exposure to a 4 Gy dose of ionizing radiation (Figure 3(B3, B4, C3, C4)) resulted in early signs of injury, including reduced neurite density and mild fragmentation of βTubulin III+ neuronal processes, accompanied by decreased OLIG1+ OPC staining. More pronounced effects were observed at the 8 Gy dose of ionizing radiation (Figure 3(B5, B6, C5, C6)), where neuronal networks appeared sparse and disorganized, OPC populations were markedly reduced, and astrocytic morphology was altered. At the highest dose tested (12 Gy; Figure 3(B7, B8, C7, C8)), cultures demonstrated substantial loss of neuronal branching, reduction in OPCs, and increased nuclear condensation as revealed by Hoechst staining, consistent with radiation-induced cellular damage. These radiation-associated phenotypes were consistently observed across both LMNSC01 and LMNSC02 lines, demonstrating reproducible and dose-responsive neural injury within the 3D LMNSC01 and LMNSC02 cultures.

### 3.4. LMNSC01-EVs and LMNSC02-EVs Promote Recovery of Neuronal Populations After MTX Exposure

To assess the restorative potential of NSC-derived EVs, differentiated LMNSC01 and LMNSC02 3D neural cultures were exposed to MTX followed by treatment with LMNSC-EVs (Figure 4A). After 7 days of differentiation in MC:NC-diff (1:1) medium, 3D cultures were treated with MTX (2 µM). Cultures were then fixed and immunostained for βTubulin III 7 days after MTX exposure. Treatment with MTX resulted in marked disruption of neuronal complexity, indicated by severe reduction in βTubulin III+ neuronal processes in both LMNSC01 (Figures 4B, 4D, S1A) and LMNSC02 (Figures 4C, 4E, S1B) 3D cultures, consistent with chemotherapy-induced neuronal injury.

**Figure 4.**
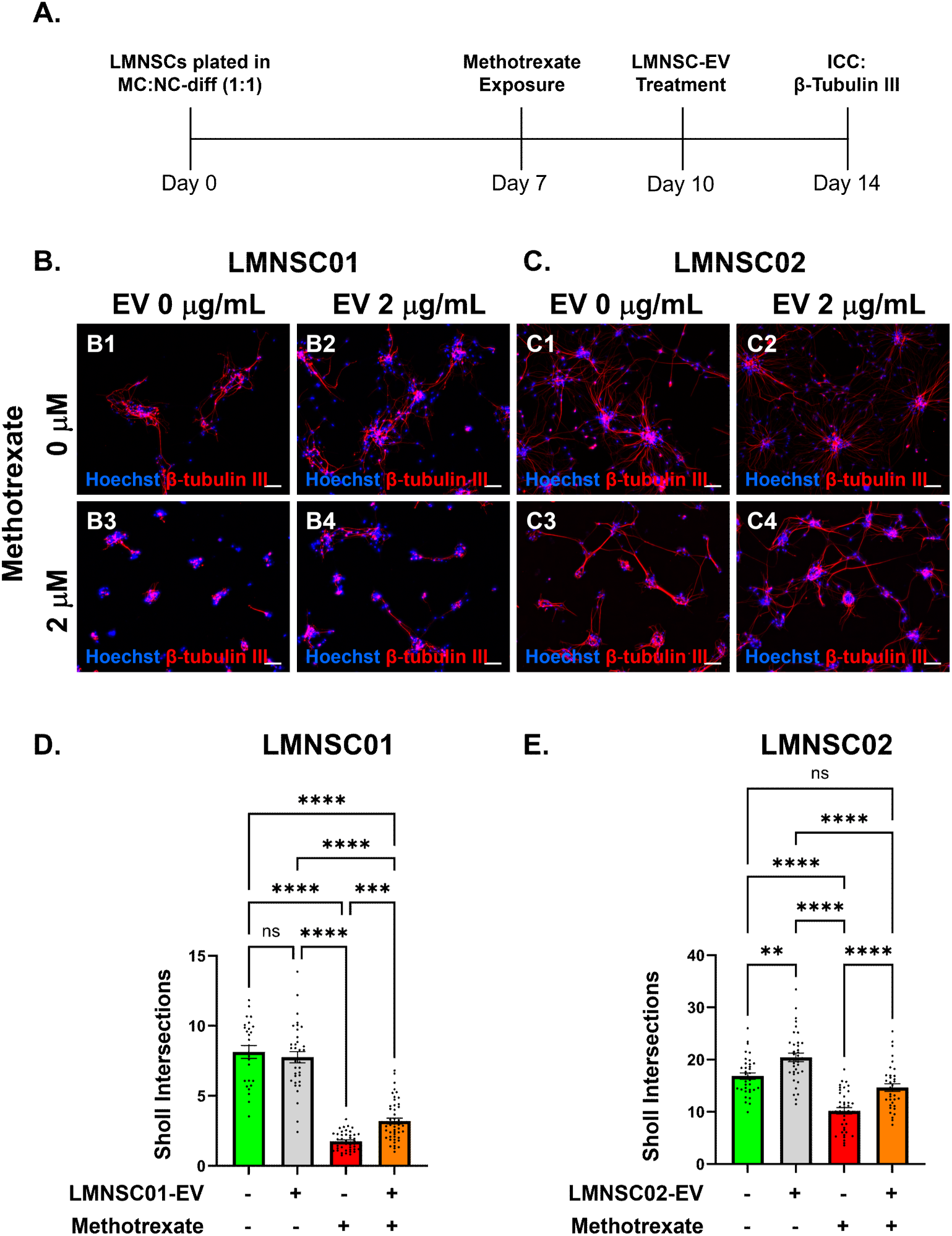
LMNSC01-EVs and LMNSC02-EVs promote recovery of neuronal populations after MTX exposure. (**A**) Experimental timeline; (**B**) representative images of ICC of LMNSC01 cells differentiated for 7 days, exposed to 2 mM MTX for 3 days, and treated with LMNSC01-EVs (2 mg/mL) for 4 days in a 3D MC:NC-diff (1:1) culture; (**C**) representative images of ICC of LMNSC02 cells differentiated for 7 days, exposed to 2 mM MTX for 3 days, and treated with LMNSC02-EVs (2mg/mL) for 4 days in a 3D MC:NC-diff (1:1) culture; (**D**) Quantification of LMNSC01 cell aggregate complexity by Sholl analysis after 14 days in 3D MC:NC-diff (1:1) culture. Each data point represents the average number of Sholl intersections of a single LMNSC01 cell aggregate. (**E**) Quantification of LMNSC02 cell aggregate complexity by Sholl after 14 days in 3D MC:NC-diff (1:1) culture. Each data point represents the average number of Sholl intersections of a single LMNSC02 cell aggregate. ICC staining was done with β-Tubulin III (red) and nuclei were stained with Hoechst (blue). Images were obtained on a Zeiss Observer II (10x). All images are representative maximum intensity projections from z-stack fluorescent images. Maximum projection and background subtraction on images were performed using Zeiss ZEN (ZEN, VER 3.12). Scale bar = 100 µm. Bars on graphs represent mean ± SEM. Results shown are from two independent experiments, where the first experiment contained one biological replicate with one technical replicate, while the second experiment contained three biological replicates with three technical replicates each. Data were analyzed using oneway ANOVA, where statistical significance was depicted as follows: ns = not significant, ** *p* ≤ 0.01, *** *p* ≤ 0.001, **** *p* ≤ 0.0001.

Cultures in the treatment group received 2 µg/mL of line-matched LMNSC-EVs 3 days after MTX exposure and were allowed to recover for an additional 4 days (Figure 4A). LMNSC-EV treatment promoted recovery of neuronal populations after MTX exposure, exhibited by increased abundance of βTubulin III+ neurons, enhanced neurite outgrowth, and, in the case of LMNSC01-EV treatment, partial reestablishment of neuronal complexity compared with MTX-treated cultures that did not receive EVs (Figures 4B, 4D, S1A). Notably, EV-mediated recovery restored neuronal complexity in LMNSC02 cultures to baseline levels (Figures 4C, 4E, S1B). These findings demonstrate that LMNSC-EVs, especially LMNSC02-EVs, support neuronal recovery and structural restoration following chemotherapy-induced injury in a 3D human neural tissue model.

### 3.5. LMNSC01-EVs and LMNSC02-EVs Promote Recovery of Glial Populations After MTX Exposure

To evaluate the effects of LMNSC-EVs on glial recovery following chemotherapy-induced injury, differentiated LMNSC01 and LMNSC02 3D neural cultures were exposed to MTX and subsequently treated with LMNSC-EVs (Figure 5A). After 7 days of differentiation in MC:NC-diff (1:1) medium, 3D cultures were treated with MTX (2 µM). Cultures were fixed and immunostained for OLIG1 and glial fibrillary acidic protein (GFAP) 7 days after MTX exposure. Treatment with MTX resulted in disruption of glial populations (Figure 5B, 5C). MTX exposure led to a significant reduction in total and OLIG1+ OPC cell number in LMNSC01 cultures (Figure 5D), and a reduction, albeit not statistically significant, in total and OLIG+ OPC cell number in LMNSC02 cultures (Figure 5E). MTX exposure also altered GFAP+ astrocyte organization (Figure 5B, 5C), with increased nuclear condensation evident by Hoechst staining, consistent with glial injury.

**Figure 5.**
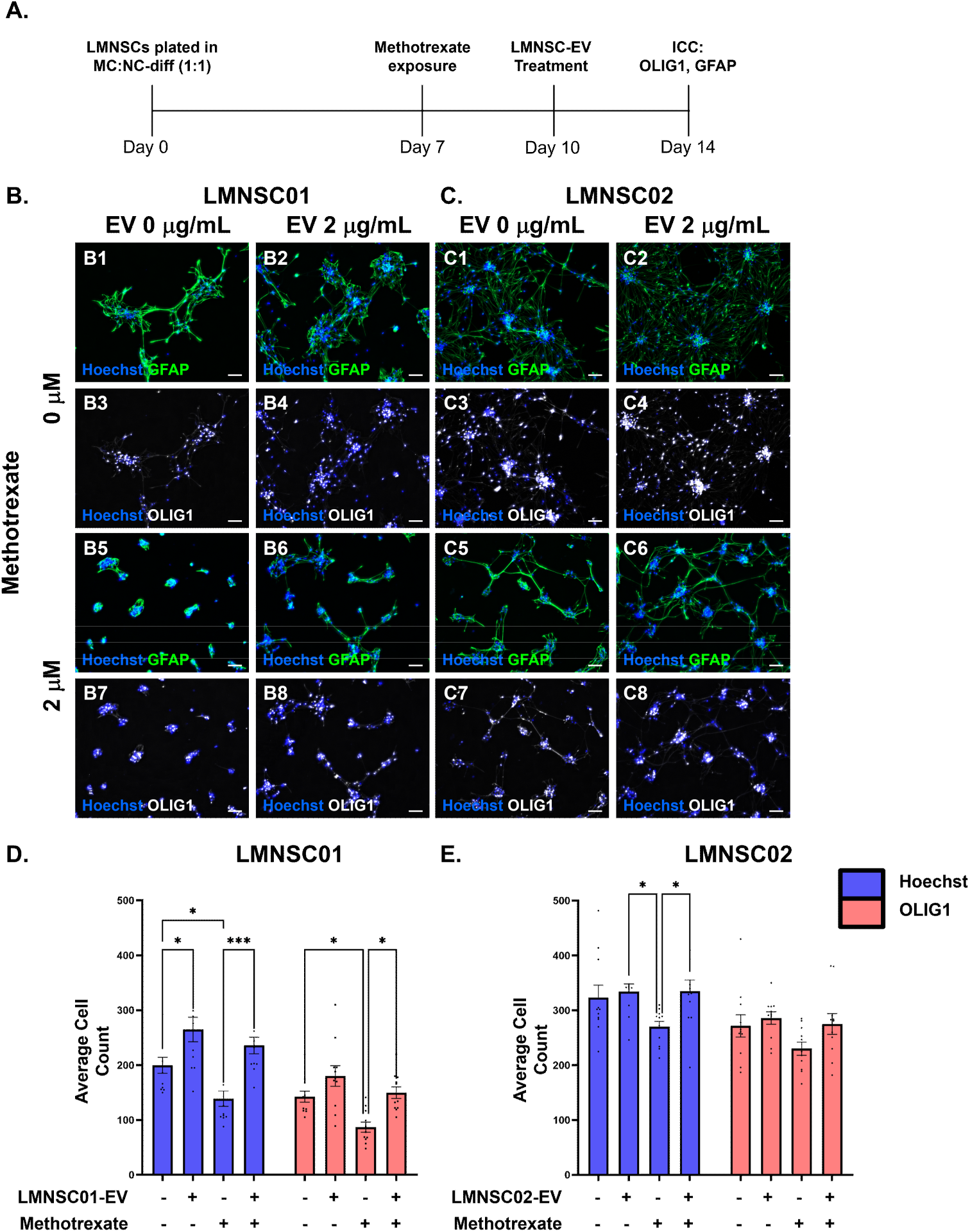
LMNSC01-EVs and LMNSC02-EVs promote recovery of glial populations after MTX exposure. (**A**) Experimental timeline; (**B**) representative images of immunocytochemistry of LMNSC01 cells differentiated for 7 days, exposed to 2 µM MTX for 3 days, and treated with LMNSC01-EVs (2 mg/mL) for 4 days in a 3D MC:NC-diff (1:1) culture; (**C**) representative images of ICC of LMNSC02 cells differentiated for 7 days, exposed to 2 mM MTX for 3 days, and treated with LMNSC02-EVs (2 mg/mL) for 4 days in a 3D MC:NC-diff (1:1) culture; (**D**) quantification of total and OLIG1+ oligodendrocyte LMNSC01 cell number after 14 days in 3D MC:NC-diff (1:1) culture. Each data point represents the cell count in a single image across two independent experiments. (**E**) Quantification of total and OLIG1+ oligodendrocyte LMNSC02 cell number after 14 days in 3D MC:NC-diff (1:1) culture. Each data point represents the cell count in a single image across two independent experiments. ICC staining was done with GFAP (green) and OLIG1 (white), and nuclei were stained with Hoechst dye (blue). Images were obtained on a Zeiss Observer II (10x). All images are representative maximum intensity projections from z-stack fluorescent images. Maximum projection and background subtraction on images were performed using Zeiss ZEN (ZEN, VER 3.12). Scale bar = 100 µm. Bars on graphs represent mean ± SEM. Results shown are from two independent experiments, where the first experiment contained two biological replicates with one technical replicate, while the second experiment contained three biological replicates with three technical replicates each. Data were analyzed using two-way ANOVA, where statistical significance was depicted as follows: * *p* ≤ 0.05, *** *p* ≤ 0.001.

Cultures in the treatment group received 2 µg/mL of line-matched LMNSC-EVs 3 days after MTX exposure and were allowed to recover for an additional 4 days (Figure 5A). LMNSC-EV treatment promoted recovery of glial populations in both LMNSC lines. Notably, treatment with LMNSC-EVs following MTX exposure restored total and OLIG1+ OPC cell numbers (Figure 5D, 5E). LMNSC-EV treatment also resulted in partial restoration of GFAP+ astrocytic networks compared with MTX-exposed cultures without EV treatment. Notably, glial recovery was observed across MTX doses, indicating that LMNSC-EVs support restoration of both oligodendroglial and astrocytic populations following chemotherapy-induced damage. These findings suggest that NSC-derived EVs facilitate glial repair and contribute to tissue recovery in a 3D human neural model of chemotherapy-associated neurotoxicity.

### 3.6. LMNSC01-EVs and LMNSC02-EVs Promote Recovery of Neuronal Populations After Irradiation

To determine whether LMNSC-EVs support neuronal recovery following radiation-induced injury, differentiated LMNSC01 and LMNSC02 3D neural cultures were exposed to ionizing radiation and subsequently treated with LMNSC-EVs (Figure 6A). After 8 days of differentiation in MC:NC-diff (1:1) medium, neural tissues were irradiated with 8 Gy dose of ionizing radiation. Cultures were then fixed and immunostained for β-Tubulin III and Hoechst 4 days after irradiation. Radiation exposure resulted in disruption of neuronal complexity, as evidenced by reduced β-Tubulin III+ neuronal density and fragmentation of neurite networks. LMNSC02 cultures exhibited a significant reduction in aggregate complexity (Figures 6E, S1D). In LMNSC01 cultures, aggregates were highly compact and exhibited a low degree of complexity. While we did not observe a reduction in aggregate complexity after irradiation, we did observe a reduced capacity of LMNSC01 cells to form aggregates, with only a total of 13 aggregates identified (Figures 6D, S1C).

**Figure 6.**
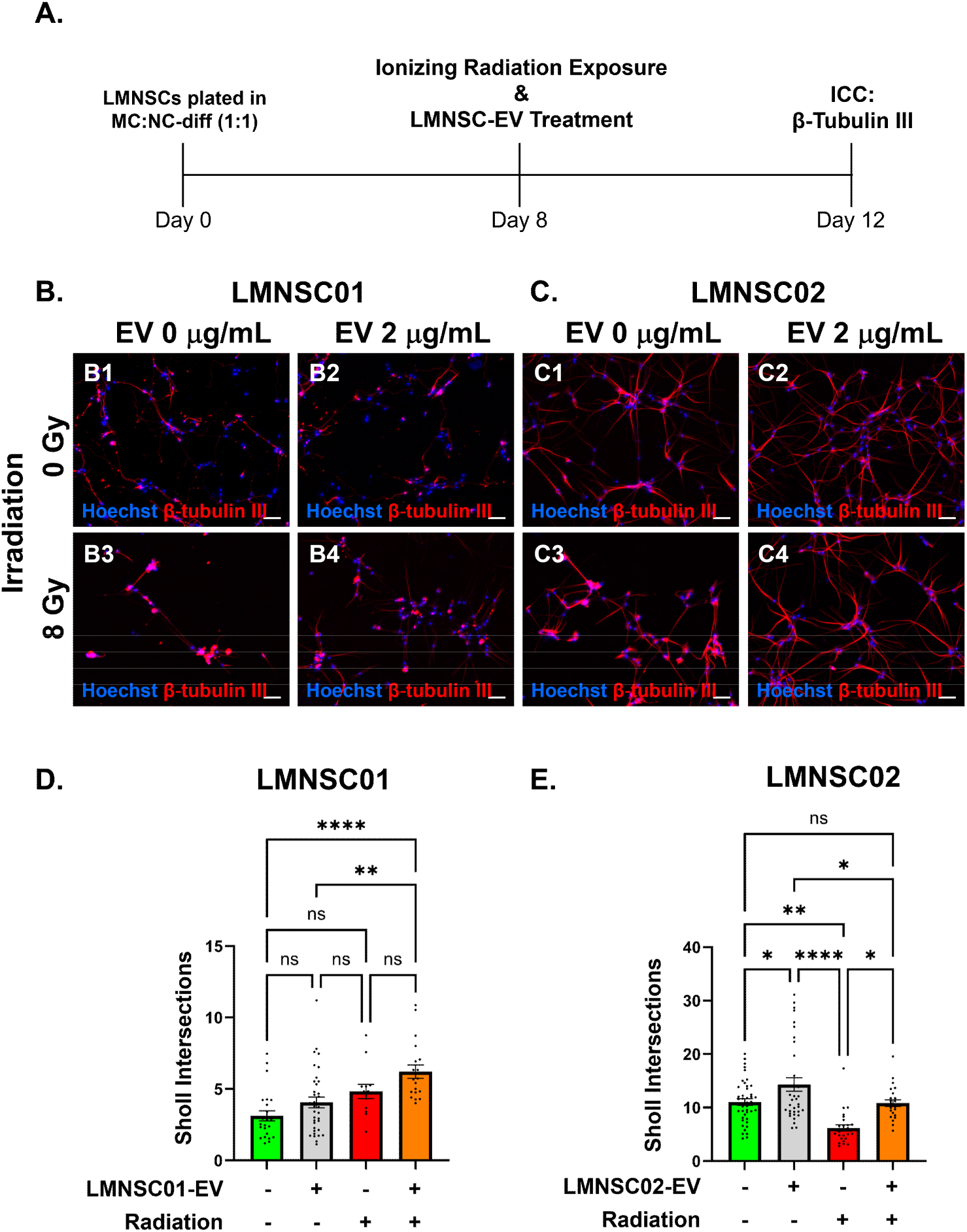
LMNSC01-EVs and LMNSC02-EVs promote recovery of neuronal populations after irradiation. (**A**) Experimental timeline; (**B**) representative images of immunocytochemistry (ICC) of LMNSC01 cells differentiated for 8 days, exposed to 8 Gy ionizing radiation, and treated with LMNSC01-EVs (2 mg/mL) for 4 days in a 3D MC:NC-diff (1:1) culture; (**C**) representative images of ICC of LMNSC02 cells differentiated for 8 days, exposed to 8 Gy ionizing radiation, and treated with LMNSC02-EVs (2 mg/mL) for 4 days in a 3D MC:NC-diff (1:1) culture; (**D**) quantification of LMNSC01 cell aggregate complexity by Sholl analysis after 12 days in 3D MC:NC-diff (1:1) culture. Each data point represents the average number of Sholl intersections of a single LMNSC01 cell aggregate. (**E**) Quantification of LMNSC02 cell aggregate complexity by Sholl analysis after 12 days in 3D MC:NC-diff (1:1) culture. Each data point represents the average number of Sholl intersections of a single LMNSC02 cell aggregate. ICC staining was done with β-Tubulin III (red), and nuclei were stained with Hoechst (blue). Images were obtained on a Zeiss Observer II (10x). All images are representative maximum intensity projections from z-stack fluorescent images. Maximum projection and background subtraction on images were performed using Zeiss ZEN (ZEN, VER 3.12). Scale bar = 100 µm. Bars on graphs represent mean ± SEM. Results shown are from two independent experiments, where the first experiment contained one biological replicate with one technical replicate, while the second experiment contained three biological replicates with three technical replicates each. Data were analyzed using one-way ANOVA, where statistical significance was depicted as follows: ns = not significant, * *p* ≤ 0.05, ** *p* ≤ 0.01, **** *p* ≤ 0.0001.

Treatment groups received 2 µg/mL of line-matched LMNSC-EVs immediately after irradiation and were allowed to recover for 4 days (Figure 6A). LMNSC-EVs treatment following irradiation promoted recovery of neuronal populations. LMNSC02-EV-treated cultures exhibited increased β-Tubulin III+ neuronal staining and enhanced neurite outgrowth compared with irradiated cultures without EV treatment. Notably, aggregate complexity was restored to baseline levels in LMNSC02 cultures (Figures 6E, S1D), indicating partial restoration of neuronal structure and complexity. These findings demonstrate that LMNSC02-EVs support neuronal recovery after radiation-induced injury in a 3D human neural tissue model.

### 3.7. LMNSC01-EVs and LMNSC02-EVs Promote Recovery of Glial Populations After Irradiation

To examine the effects of LMNSC-EVs on glial recovery following radiation-induced injury, differentiated LMNSC01 and LMNSC02 3D neural cultures were exposed to ionizing radiation and subsequently treated with LMNSC-EVs (Figure 7A). After 8 days of culture in MC:NC-diff (1:1) medium, neural tissues were irradiated with an 8 Gy dose of ionizing radiation. Cultures were then fixed and immunostained for OLIG1, GFAP, and Hoechst 4 days after irradiation. Exposure to ionizing radiation resulted in pronounced glial injury (Figure 7B, 7C). Irradiated cultures exhibited a significant reduction in total and OLIG1+ OPC cell number in both LMNSC01 (Figure 7D) and LMNSC02 (Figure 7E) cultures. Additionally, we observed disruption of GFAP+ astrocytic networks, accompanied by increased nuclear condensation detected by Hoechst staining, indicating radiation-induced glial damage.

**Figure 7.**
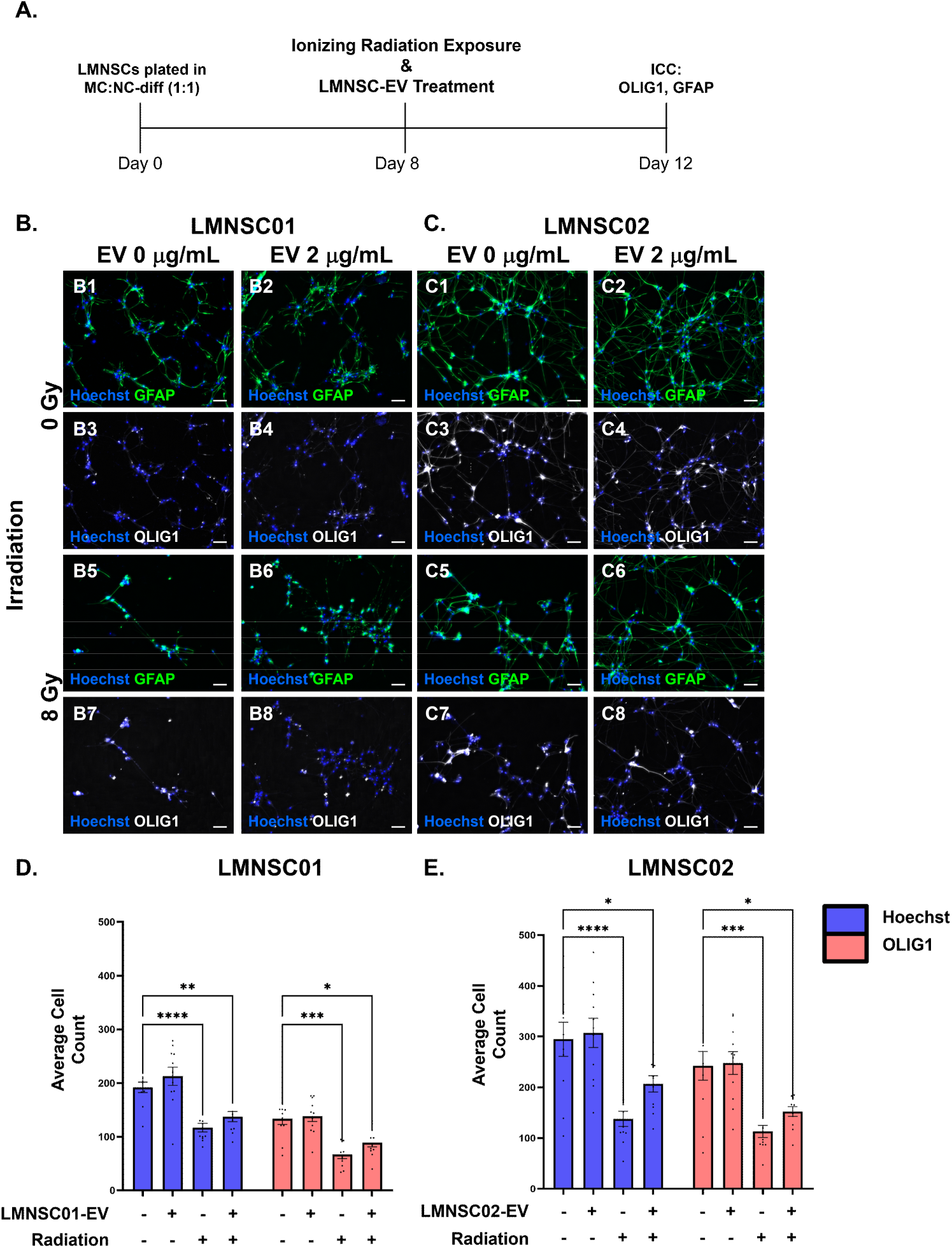
LMNSC01-EVs and LMNSC02-EVs promote recovery of glial populations after irradiation. (**A**) Experimental timeline; (**B**) representative images of immunocytochemistry of LMNSC01 cells differentiated for 8 days, exposed 8 Gy ionizing radiation, and treated with LMNSC01-EVs (2 mg/mL) for 4 days in a 3D MC:NC-diff (1:1) culture; (**C**) representative images of ICC of LMNSC02 cells differentiated for 8 days, exposed to 8 Gy ionizing radiation, and treated with LMNSC02-EVs (2 mg/mL) for 4 days in a 3D MC:NC-diff (1:1) culture; (**D**) quantification of total and OLIG1+ oligodendrocyte LMNSC01 cell number after 12 days in 3D MC:NC-diff (1:1) culture. Each data point represents the cell count in a single image across two independent experiments. (**E**) Quantification of total and OLIG1+ oligodendrocyte LMNSC02 cell number after 12 days in 3D MC:NC-diff (1:1) culture. Each data point represents the cell count in a single image across two independent experiments. ICC staining was done with GFAP (green) and OLIG1 (white), and nuclei were stained with Hoechst dye (blue). Images were obtained on the Zeiss Observer II (10x). All images are representative maximum intensity projections from z-stack fluorescent images. Maximum projection and background subtraction on images were performed using Zeiss ZEN (ZEN, VER 3.12). Scale bar = 100 µm. Bars on graphs represent mean ± SEM. Results shown are from two independent experiments, where the first experiment contained two biological replicates with one technical replicate, while the second experiment contained three biological replicates with three technical replicates each. Data were analyzed using two-way ANOVA, where statistical significance was depicted as follows: * *p* ≤ 0.05, ** *p* ≤ 0.01, *** *p* ≤ 0.001, **** *p* ≤ 0.0001.

Treatment with 2 µg/ml of line-matched LMNSC-EVs for 4 days following irradiation promoted partial recovery of glial populations in both LMNSC cultures. LMNSC-EV-treated cultures demonstrated increased total and OLIG1+ OPC cell numbers in both LMNSC01 (Figure 7D) and LMNSC02 (Figure 7E) cultures, although not restored to levels comparable to those of control groups, indicating that LMNSC-EVs support recovery of both total and OLIG1+ OPC cell numbers following irradiation. Additionally, treatment with respective LMNSC-EVs promoted recovery of GFAP+ astrocytic networks in both LMNSC cultures (Figure 7B, 7C), indicating that LMNSC-EVs support repair of astrocytic populations following radiation-induced injury. These results demonstrate that NSC-derived EVs facilitate glial restoration in a 3D human neural tissue model of radiation-associated neurotoxicity.

### 3.8. Radiation-Induced Transcriptional Programs Are Reversed by LMNSC01-EV Treatment

To evaluate the effects of LMNSC-EV treatment following radiation-induced injury on the transcriptome, differentiated LMNSC01 cultures were exposed to ionizing radiation and subsequently treated with LMNSC01-EVs (Figure 8A). After 8 days of differentiation, LMNSC01 cells were irradiated with an 8 Gy dose of ionizing radiation. Bulk RNAseq was performed 4 days after exposure to 8 Gy radiation. Differential expression analysis identified significant alterations in transcriptional profile in irradiated cultures compared to untreated controls, including 1678 upregulated and 2121 downregulated genes (Figure 8B). Among the 10 most significant differentially expressed genes (DEGs), several neural lineage-associated transcripts were markedly downregulated following irradiation, including SVOP (log2FC = −4.176), NSG2 (log2FC = −3.927), GPR139 (log2FC = −4.176), NEUROG2 (log2FC = −4.176), and SCGN (log2FC = −4.176) (Figure 8C). Conversely, genes associated with stress response were found to be significantly upregulated, including CAVIN3 (log2FC = 3.977) and TMTB1 (log2FC = 3.785). These results suggest that exposure to ionizing radiation reduces the expression of genes that are associated with neuronal lineage specification and differentiation while also inducing the expression of genes involved in stress response, in line with neural injury.

**Figure 8.**
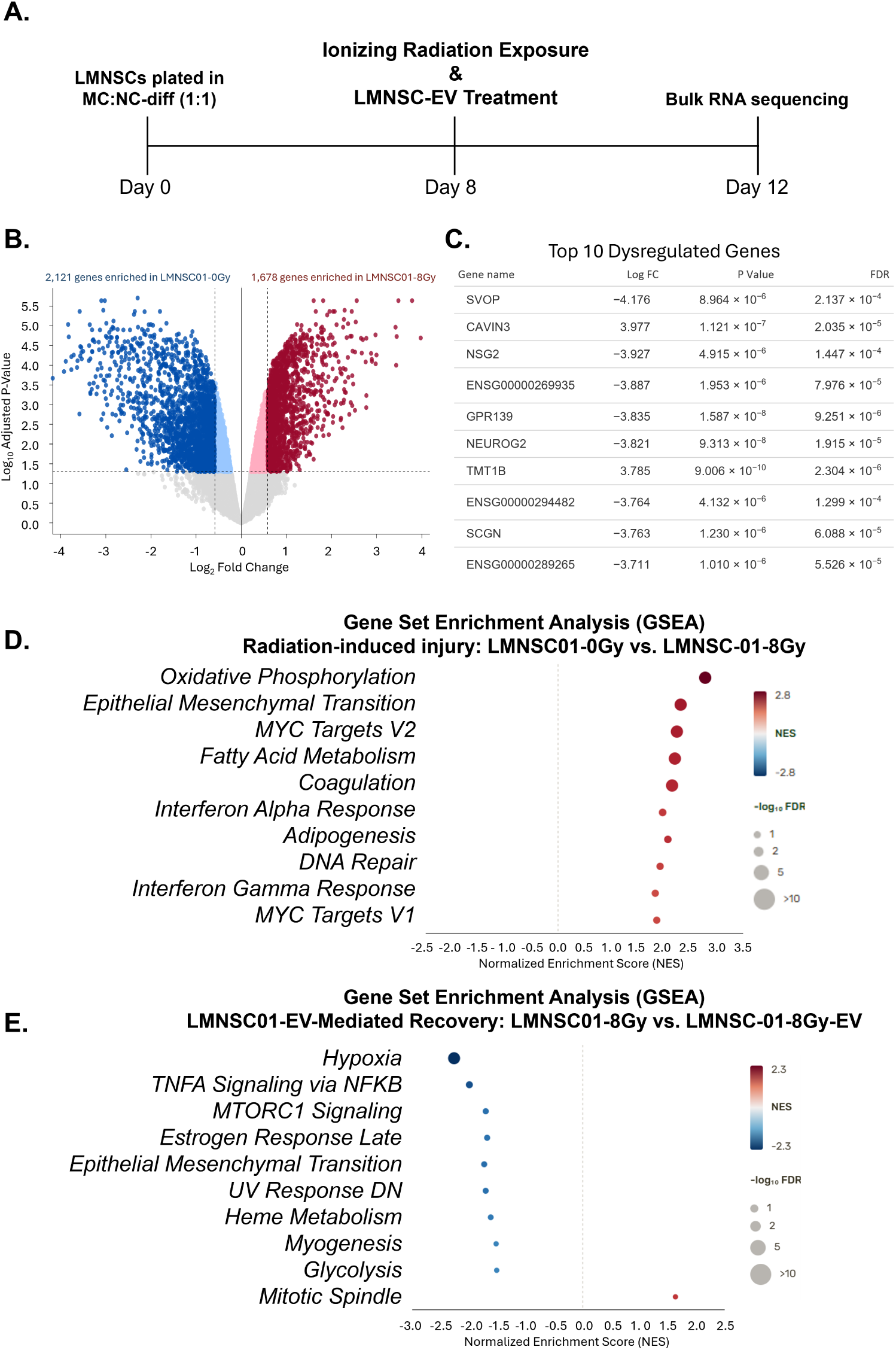
Radiation-induced transcriptional programs are reversed by LMNSC01-EV treatment. (**A**) Experimental timeline and (**B**) volcano plot showing DEGs in irradiated LMNSC01 cultures compared to untreated controls. Red and blue points indicate significantly upregulated (log₂FC ≥ 0.585, *p* < 0.05) and downregulated (log₂FC ≤ −0.585, *p* < 0.05) genes, respectively. (**C**) List of the top 10 DEGs in irradiated LMNSC01 cultures compared to untreated controls. (**D**) Gene set enrichment analysis (GSEA) showing enriched pathways in irradiated LMNSC01 cultures compared to untreated controls, analyzed with the MSigDB Hallmarkgene set. Pathways were sorted by adjusted *p*-value in ascending order. (**E**) Gene set enrichment analysis (GSEA) showing enriched pathways in irradiated LMNSC01 cultures treated with LMNSC01-EVs compared to irradiated LMNSC01 cultures analyzed with the MSigDB Hallmarkgene set. Pathways were sorted by adjusted *p*-value in ascending order.

Gene set enrichment analysis (GSEA) of irradiated cultures highlighted significant enrichment of pathways associated with oxidative phosphorylation, epithelial–mesenchymal transition, MYC targets, fatty acid metabolism, interferon-α and -γ response, and DNA repair, amongst others (Figure 8D). Enrichment of genes involved in DNA repair and interferon pathways indicates activation of cellular stress and damage response mechanisms. The high normalized enrichment score (NES) of oxidative phosphorylation and fatty acid metabolism suggests that a metabolic shift occurs in response to irradiation to support high energy demands involved in stress and damage response.

To determine the ability of LMNSC01-EVs to modulate the radiation-induced transcriptional response, GSEA was performed on EV-treated irradiated cultures compared to irradiated cultures alone. LMNSC01-EV treatment resulted in negative NES for hypoxia, TNFα signaling via NFκB, MTORC1, epithelial–mesenchymal transition, UV response, and glycolysis pathways, among others (Figure 8E). In contrast, a positive NES was observed for the mitotic spindle pathway. These findings suggest that LMNSC01-EVs reverse radiation-induced stress, inflammatory response, and metabolic normalization, as evidenced by inhibition of hypoxia, TNFα signaling via NFκB, UV response, and epithelial–mesenchymal transition signaling pathways. Furthermore, the activation of the mitotic spindle pathway suggests that LMNSC01-EV treatment promotes cell-cycle progression, supporting the idea that they can promote recovery of proliferative activity in irradiated LMNSCs.

Collectively, these findings demonstrate that LMNSC01-EVs broadly reverse radiation-induced transcriptional reprogramming by attenuating inflammatory, stress, and metabolic injury responses while promoting pathways associated with cellular recovery and regeneration. Rather than targeting a single pathway, LMNSC01-EVs orchestrate co-ordinated remodeling of multiple biological processes that underlie radiation-induced neural injury.

## 4. Discussion

In this study, we developed a human NSC-derived 3D neural tissue model generated from LMNSCs that provides an experimentally tractable platform for studying chemotherapy- and radiation-induced neural injury. Using this model, we investigated the effects of cancer therapies on neuronal, astrocytic, and oligodendroglial populations and evaluated the neuroprotective and regenerative potential of NSC-derived EVs. Our findings demonstrate that LMNSC-derived 3D neural cultures generate multicellular neural tissues containing neuronal, astrocytic, and oligodendroglial populations that resemble key components of the central nervous system. These cultures provide a platform in which neuronal, astrocytic, and oligodendrocyte lineage cells interact within a structured microenvironment. Such models address important limitations of conventional two-dimensional neural cultures, which lack tissue architecture and complex cellular interactions, and complement animal models that may not fully reflect human-specific responses to therapy. The ability of this platform to generate organized neural tissues suggests that it can serve as a valuable experimental system for studying therapy-induced neurotoxicity and neural repair.

Using this model, we demonstrated that exposure to chemotherapy and ionizing radiation induces significant neural injury characterized by loss of neuronal and oligodendrocyte progenitor populations, and that ionizing radiation induces alterations in transcriptional pathways associated with neuroinflammation and cellular stress. These findings are consistent with previous studies showing that chemotherapeutic agents such as MTX impair neurogenesis and damage OPCs, leading to disruptions in white matter integrity and neuronal signaling [8]. Radiation exposure has similarly been shown to disrupt NSC niches and promote chronic inflammatory responses that contribute to longterm neurodegeneration [33]. Our results extend these observations by demonstrating that therapy-induced injury can be reproduced in human NSC-derived 3D cultures, providing a controlled platform for mechanistic investigation.

Importantly, our study also demonstrates the therapeutic potential of NSC-derived EVs in mitigating cancer therapy-induced neural damage. Interestingly, both LMNSC lines exhibited distinct responses to MTX-induced injury and EV-mediated recovery [25]. LMNSC01 cultures demonstrated increased sensitivity to MTX exposure, evident by a significant reduction in OPC populations, whereas LMNSC02 cultures showed comparatively less substantial oligodendroglial loss. Notably, treatment with matched LMNSC01-EVs restored OPC numbers to levels comparable to untreated controls, suggesting a potent effect on oligodendroglial recovery. In contrast, although LMNSC02-EVs also promoted oligodendroglial recovery, their most pronounced effects were observed in the restoration of neuronal morphology and neurite complexity. These findings suggest that intrinsic differences between LMNSC lines may influence both susceptibility to therapy-induced injury and the regenerative properties of their corresponding EVs. These findings support growing evidence that many of the beneficial effects of stem cell therapies are mediated through paracrine mechanisms, particularly through EVs that deliver bioactive molecules to recipient cells [30,34]. EVs carry a diverse cargo of proteins, lipids, and regulatory RNAs capable of modulating cellular signaling pathways involved in inflammation, synaptic plasticity, and tissue repair. In the context of therapy-induced brain injury, EV-mediated signaling may therefore represent a promising cell-free strategy for promoting neural recovery while avoiding some of the challenges associated with direct stem cell transplantation.

Our transcriptomic analyses demonstrate that radiation induces extensive molecular reprogramming in differentiating LMNSC cultures, characterized by suppression of neuronal differentiation programs and activation of inflammatory, metabolic, and DNA damage-response pathways. These changes are consistent with persistent cellular stress and impaired differentiative capacity that contribute to radiation-induced neurotoxicity. It is worth noting that our 3D neural tissue model lacks the primary immune mediator of the brain, microglia. As such, the inflammatory response pathways identified here should therefore not be interpreted as representing the full neuroimmune response to irradiation or the microglia-dependent mechanisms that are implicated in CRCI [8,35–37]. Rather, these signatures reflect the inflammatory and stress-associated transcriptional responses intrinsic to the LMNSC-derived neural populations within the cultures. Treatment with LMNSC01-EVs was associated with altered enrichment of several injury-associated transcriptional programs, suppressing hypoxia, TNFα/NF-κB signaling, epithelial–mesenchymal transition, mTORC1 signaling, glycolysis, and other stress-response pathways while promoting mitotic spindle signaling. These findings suggest that LMNSC01-EVs act as pleiotropic biological regulators of signaling pathways associated with inflammation, cellular metabolism, and tissue repair rather than targeting a single molecular pathway. The coordinated attenuation of multiple radiation-associated transcriptional networks supports the therapeutic potential of LMNSC01-EVs as a regenerative strategy to mitigate radiation-induced neural injury and preserve brain function following cranial irradiation.

The use of LMNSC-EV-based therapies may be particularly advantageous in oncology settings where safety considerations are paramount. While stem cell transplantation has shown promise in regenerative medicine, concerns regarding engraftment, immune responses, and potential carcinogenic effects remain important considerations. The use of EVs may circumvent some of these limitations by delivering therapeutic signals without the risks associated with cellular therapies. Previous studies have demonstrated that EV-based interventions can promote neuroprotection, reduce injury-associated inflammatory responses, improve neuronal integrity, and support functional recovery across a variety of preclinical models of neurological injury, including radiation-induced CRCI and neurodegenerative diseases [27,38–40]. Our findings add to this growing body of evidence by demonstrating LMNSC-EV-associated recovery of neuronal, astrocytic, and oligodendroglial lineage populations, as well as attenuation of injury-associated transcriptional signatures, within a 3D human LMNSC-based neural tissue model. While the absence of microglia limits interpretation of microglia-dependent neuroinflammatory mechanisms, these findings provide insight into direct cellular responses of human neural-lineage populations following injury and EV treatment. In addition, LMNSC-EVs can be engineered or enriched for specific cargo to enhance their neuroprotective and neurorestorative effects, offering opportunities for precision therapeutic strategies. Characterization of LMNSC01-EVs and LMNSC02-EVs demonstrated significant differences in EV size, tetraspanin expression, and protein cargo [25]. These observations support the use of donor-matched EVs in the present study and suggest that donor-specific EV characteristics may influence their therapeutic efficacy. Future studies should evaluate whether cross-donor EV delivery produces comparable neuroprotective effects, which would support the potential use of LMNSC-EVs as an allogeneic product for treating neural injury.

Beyond the therapeutic implications of LMNSC-EVs, the LMNSC-based 3D neural tissue platform described in this study provides a versatile tool for modeling treatment-related neurotoxicity in a human-relevant system. Recently, we characterized LMNSC01 and LMNSC02 cells by single-cell transcriptomics and demonstrated that both lines represent distinct NSC populations with minimal transcriptional overlap and divergent differentiation trajectories [25]. This is in line with the phenotypic differences we observed between the LMNSC01- and LMNSC02-based 3D neural tissues in this study, suggesting that these differences are primarily attributable to intrinsic donor-specific characteristics rather than culture-induced variation. While these donor-specific characteristics introduce biological variability, they may enhance the translational relevance of the model by better reflecting the heterogeneity observed among human patients. Such models could be used to investigate mechanisms underlying CRCI across different cancer therapies and to screen candidate neuroprotective or neurorestorative agents. The integration of transcriptomic profiling with functional analyses further enables the identification of molecular pathways involved in neural injury and recovery, providing insights that may inform the development of targeted interventions.

Several limitations of the present study should be acknowledged. Although the 3D neural cultures recapitulate key neuronal, astrocytic, and oligodendroglial populations, they do not fully reproduce the complexity of the in vivo brain microenvironment, which includes vascular components, immune cells, and systemic influences. In particular, the cultures do not contain microglia, which are established mediators of neuroinflammation. As a result, this model does not evaluate microglial activation, complement-mediated synaptic pruning, microglia-neuron interactions, or the full neuroimmune response to MTX or ionizing radiation. Furthermore, the markers evaluated in this study primarily identify neural lineage commitment rather than maturational or functional state of the cultures. Accordingly, the findings should be interpreted in the context of differentiating neural lineage populations rather than fully mature neural populations. Future studies incorporating additional cell types such as microglia or endothelial cells, together with characterizing the maturational and functional states, may enhance the physiological relevance of the model. Additionally, while our findings demonstrate EV-mediated neuroprotection at the cellular and molecular levels, further studies will be required to determine how these effects translate to functional outcomes in vivo.

Despite these limitations, the results presented here establish a human LMNSC-derived 3D neural tissue platform that can model chemotherapy- and radiation-induced brain injury and evaluate regenerative therapeutic strategies. The ability of LMNSC-EVs to mitigate therapy-induced neural damage highlights their potential as a neuroprotective intervention for cancer survivors. As survival rates continue to improve, addressing longterm neurological complications of cancer therapy has become increasingly important. Human neural tissue models combined with regenerative EV-based therapies may therefore provide a powerful framework for advancing both mechanistic understanding and therapeutic development in this emerging area of cancer survivorship research.

## 5. Conclusions

In this study, we developed a human 3D neural tissue model derived from LMNSCs to investigate mechanisms of chemotherapy- and radiation-induced neurotoxicity and to evaluate EV-mediated neuroprotection. Our results demonstrate that LMNSC-derived 3D neural tissue cultures recapitulate key neuronal, astrocytic, and oligodendroglial populations and provide a platform for modeling therapy-induced brain injury and EV-mediated recovery. Exposure to MTX and ionizing radiation induced lineage-specific neural damage and transcriptional changes associated with neuroinflammation, cellular stress, and impaired neuronal signaling. Importantly, treatment with NSC-derived EVs promoted recovery of neural populations and partially restored molecular pathways involved in neuronal survival and tissue repair. Together, these findings highlight the potential of human 3D neural tissue models for studying CRCI and support the use of NSC-derived EVs as a promising cell-free therapeutic strategy to mitigate neurotoxicity associated with cancer treatment.

## Supplementary Materials

The following supporting information can be downloaded at https://www.mdpi.com/article/doi/s1. Figure S1: LMNSC-EVs promote recovery of neuronal complexity after MTX and radiation exposure. Figure S2: Radiation induces dose-dependent cellular injury and transcriptional alterations in LMNSC-derived organoids.

## Author Contributions

**Author Contributions:** Conceptualization, L.G.A.N., M.G., S.P., and S.Y.; methodology, L.G.A.N., R.C.R., and M.G.; software, R.C.R. and B.E.; validation, B.E., I.V., and L.G.A.N.; formal analysis, L.G.A.N., I.V., B.E., and R.C.R.; investigation, L.G.A.N., I.V., B.E., and L.C.; resources, R.C.R. and M.G.; data curation, B.E. and L.G.A.N.; writing—original draft preparation, L.G.A.N., R.C.R., S.P., S.Y., and M.G.; writing—review and editing, S.P., S.Y., R.C.R., and M.G.; visualization, L.C.; supervision, M.G.; project administration, M.G.; funding acquisition, M.G. All authors have read and agreed to the published version of the manuscript.

## Institutional Review Board Statement

Not applicable.

## Informed Consent Statement

Not applicable.

## Data Availability Statement

The original data presented in this study will be openly available in **GEO** on November 1, 2026. These data include the datasets generated and analyzed during the current study and are available without restriction.

## Acknowledgments

The authors thank Erin S. Keebaugh for her expert scientific writing and editorial assistance in the preparation of this manuscript. Her contributions to improving the clarity, organization, and presentation of the manuscript are gratefully acknowledged. We also thank Brian Armstrong in the Light Microscopy Imaging Core for his assistance with image acquisition and analysis.

## Abbreviations

The following abbreviations are used in this manuscript:

3D: Three-dimensional
CRCI: Cancer therapy-related cognitive impairment
EV: Extracellular vesicle
GFAP: Glial fibrillary acidic protein
LMNSC: L-MYC immortalized neural stem cell
LMNSC-EV: L-MYC immortalized neural stem cell-derived extracellular vesicle
MTX: Methotrexate
NSC: Neural stem cell
OPCs: Oligodendrocyte precursor cells

## Disclaimer/Publisher’s Note

The statements, opinions and data contained in all publications are solely those of the individual author(s) and contributor(s) and not of MDPI and/or the editor(s). MDPI and/or the editor(s) disclaim responsibility for any injury to people or property resulting from any ideas, methods, instructions or products referred to in the content.

