## Supplemental Figures for "Extracellular Vesicles Derived from *L-MYC* Neural Stem Cells Mediate Neuroprotection in 3D Models of Chemotherapy- and Radiation-Induced Neurotoxicity"

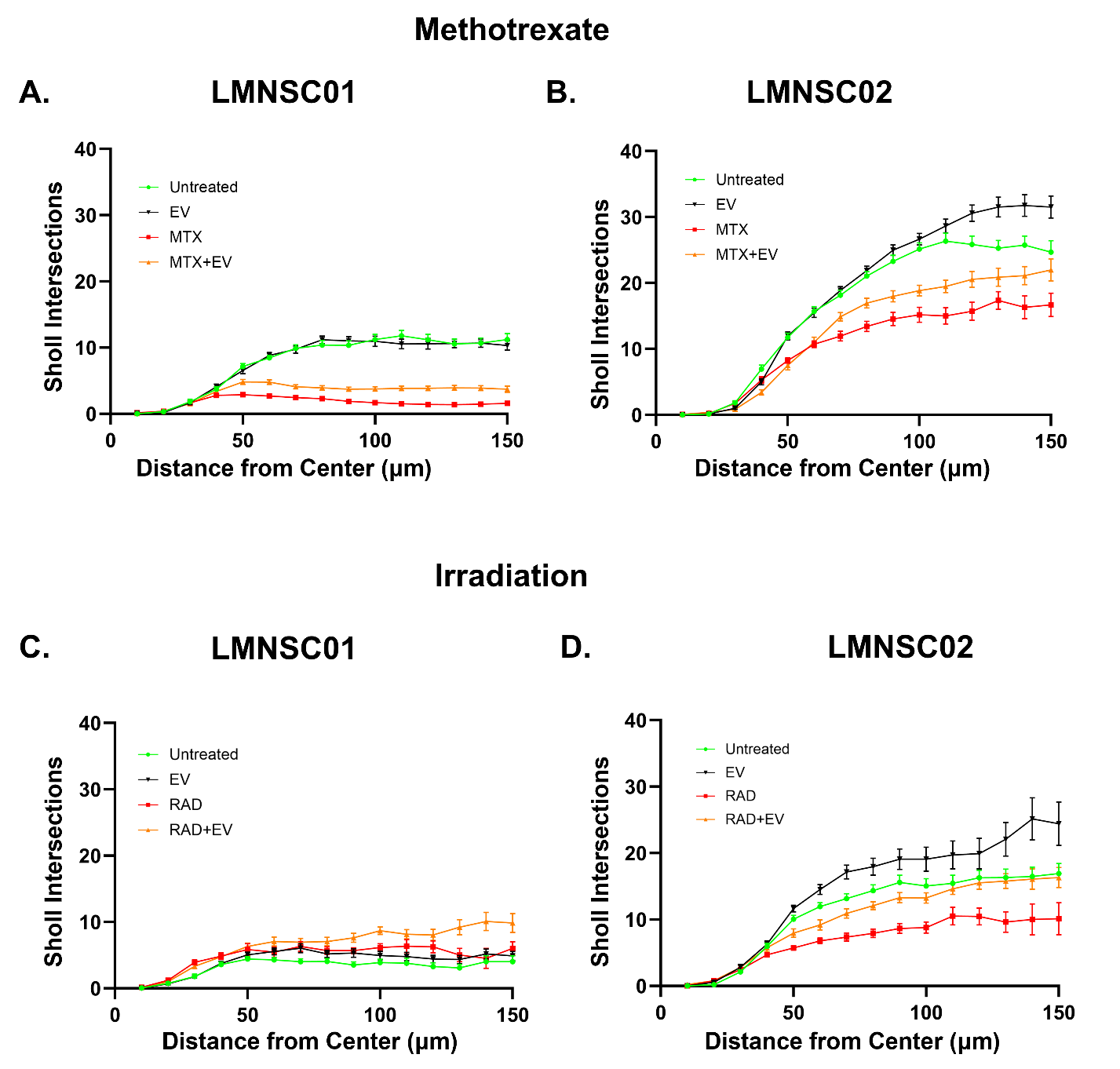
**Figure S1:  LMNSC-EVs promote recovery of neuronal complexity after methotrexate and radiation exposure.** (A) Sholl analysis intersection profile of LMNSC01 neuronal complexity, measured by βtubulin III staining, after 14 days in 3D MC:NC-diff (1:1) culture. (B) Sholl analysis intersection profile of LMNSC02 neuronal complexity, measured by βtubulin III staining, after 14 days in 3D MC:NC-diff (1:1) culture. (C) Sholl analysis intersection profile of LMNSC01 neuronal complexity, measured by βtubulin III staining, after 12 days in 3D MC:NC-diff (1:1) culture. (D) Sholl analysis intersection profile of LMNSC02 neuronal complexity, measured by βtubulin III staining, after 12 days in 3D MC:NC-diff (1:1) culture. Each datapoint represents the mean ± SEM number of Sholl intersections at increasing distances from the center of neural aggregates. Results shown are from two independent experiments, where the first experiment contained one biological replicate with one technical replicate, while the second experiment contained three biological replicates with three technical replicates each.

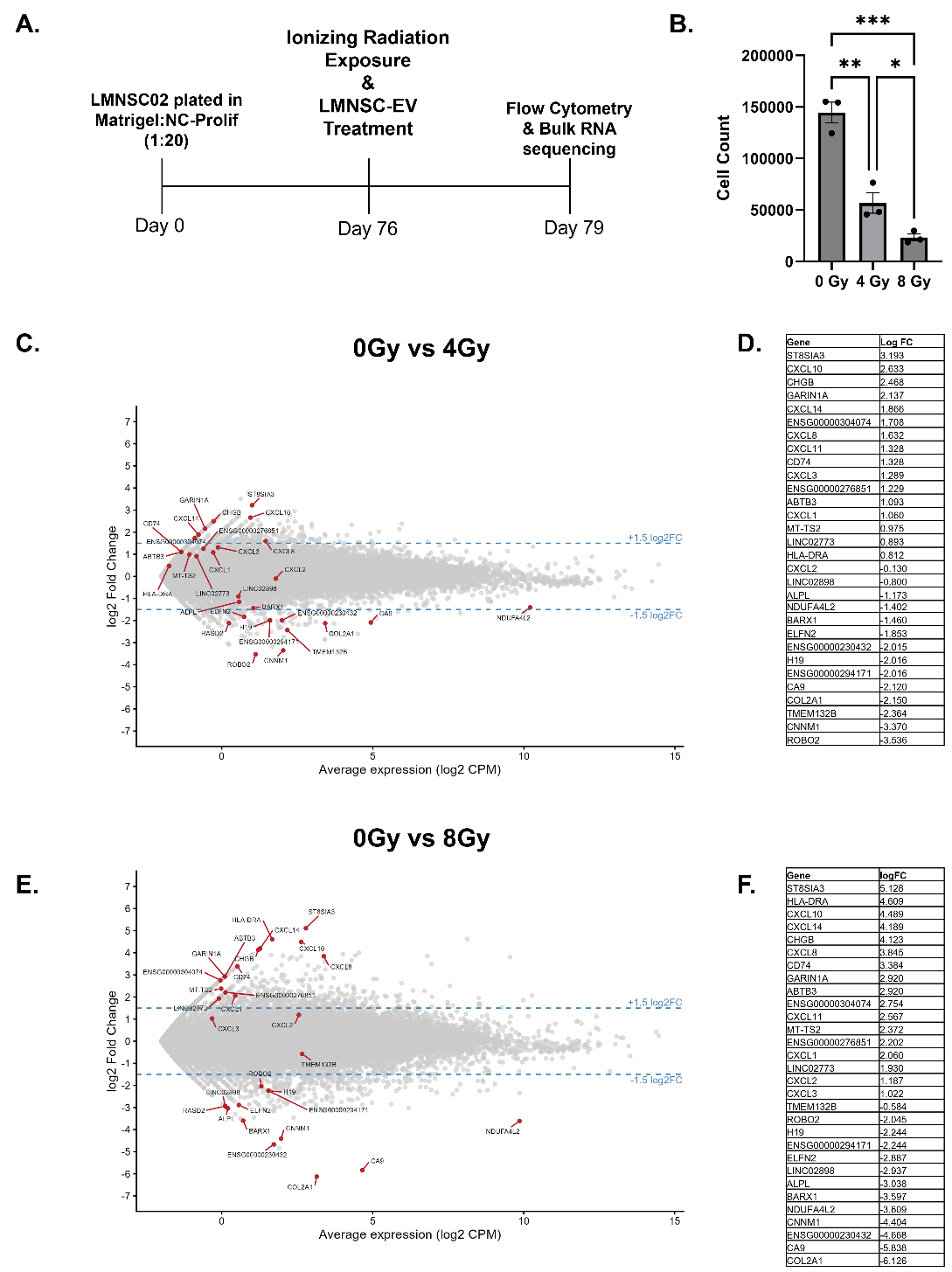
**Figure S2: Radiation induces dose-dependent cellular injury and transcriptional alterations in LMNSC-derived organoids.** (A) Experimental timeline. (B) Cell count in LMNSC02 organoids after 0, 4, and 8 Gy ionizing radiation exposure analyzed by flow cytometry. N=3, error bars represent Mean ± SEM. **p<*0.05, ***p*<0.01, ****p*<0.001. **(C)** MA plot in irradiated LMNSC02 organoids at the dose of 4Gy compared to untreated control. **(D)** List of the top 30 most differentially expressed genes in irradiated LMNSC02 organoids at the dose of 4Gy compared to untreated control. Genes were sorted by Log2 Fold Change in descending order. **(E)** MA plot in irradiated LMNSC02 organoids at the dose of 8Gy compared to untreated control. **(F)** List of the top 30 most differentially expressed genes in irradiated LMNSC02 organoids at the dose of 8Gy compared to untreated control. Genes were sorted by Log2 Fold Change in descending order.

**Supplementary Methods**

*Generation of LMNSC02 Organoid Cultures*

Organoids were generated from LMNSC02 cells. Organoids were created using a previously specified protocol [1]. Briefly, 83,000 LMNSC02 cells were suspended in 25uL of NC-prolif supplemented with EGF and bFGF. The aliquot was combined with 500uL of Matrigel (Corning, Cat#356231) and 6uL of matrigel-cell suspension was placed on previously sterilized parafilm squares in a 6 well plate (Matrigel:NC-prolif) (1:20). Freshly made organoids were incubated under physiological hypoxia (4% O2, 5% CO2) for 3 hours to allow gel to stiffen. Afterwards, parafilm was removed and resulting organoids were placed in 4mL of NC-prolif supplemented with EGF and bFGF. Organoids were transferred to orbital shaker set to 100rpm and were maintained in physiological hyopixa throughout the experiment. NC-prolif with EGF and FGF was replaced weekly for the first 3 weeks, and twice a week afterwards up until irradiation.

*Irradiation Exposure of LMNSC02 Organoids Cultures and Treatment with LMNSC-EVs*

On day 76, organoids were prepped for irradiation. An hour prior to irradiation, three organoids were added per well in a 6 well plate in 3mL of NC-prolif with 2x concentration of EGF and bFGF. Organoids were irradiated at 0Gy, 4Gy, and 8Gy using MultiRad 160 (Precision X-Ray Irradiation, Madison, US). Organoids were irradiated using the following settings: (Dose 1Gy/min, FSD 37cm, Filter: Copper 0.30mm, kV 160, mA 13.3). Immediately after irradiation, medium was aspirated and 3mL of fresh medium containing LMNSC-EVs (2 mg/mL) with EGF and bFGF was added to the wells. Organoids were left to incubate for an additional 72h to allow for recovery. Organoids were cultured in NC-prolif medium for a total of 79 days.

*RNA Isolation of LMNSC02 Organoids Cultures*

Three organoids per group were placed in a single 1.5mL tube, and homogenized with TRIzol (Invitrogen, Cat# 1559602). Organoids were disrupted and RNA was extracted using the RNeasy Mini kit (Qiagen, Cat# 74104) in accordance to the manufacturers’ protocol. Off-column DNase treatment was achieved using TURBO DNase™ (2 U/μL) (Invitrogen, Cat# AM2238). RNA clean up was performed using RNeasy MinElute Cleanup Kit (Qiagen, Cat# 74204). Due to excess phenol and guanidine contamination after RNA clean up, an ethanol precipitation step was also performed using a previously described protocol [2]. Briefly, sodium acetate (3M, 5.2 PH) was diluted to 0.3M in isolated RNA suspended in nuclease free water. 3 volumes of 100% ice cold ethanol was then added to the RNA samples and left to precipitate overnight at 4C. Supernatant was removed using centrifugation at 4C, and precipitated RNA was washed twice with 70% of ice cold ethanol. The resulting ethanol was removed and precipitated RNA pellets were suspended in nuclease free water. RNA concentration and purity was examined using Nanophotometer NP80 (Implen).

*RNA-seq Analysis of LMNSC02 Organoids*

Total RNA isolated from LMNSC02-derived brain organoids was sequenced at Plasmidsaurus (Louisville, KY). Raw base call (BCL) files were converted to FASTQ format and demultiplexed using BCL Convert (v4.3.6) and fqtk (v0.3.1). Reads were processed with FastP (v0.24.0) for poly-X trimming, 3′ quality trimming, enforcement of a minimum Phred quality score of 15, and a minimum read length of 50 bp. Filtered reads were aligned to the GRCr8v114 reference genome using STAR aligner (v2.7.11) with non-canonical splice junction removal and unmapped read output enabled. Resulting BAM files were coordinate-sorted using samtools (v1.22.1), and PCR and optical duplicates were removed using UMICollapse (v1.1.0) based on unique molecular identifiers (UMIs). Alignment quality metrics, strand specificity, and genomic feature distribution were assessed using RSeQC (v5.0.4) and Qualimap (v2.3), and a comprehensive quality control report was generated with MultiQC (v1.32). Gene-level quantification was performed using featureCounts (Subread package v2.1.1) with strand-specific counting, fractional assignment of multi-mapping reads, and annotation of exons and 3′ UTR regions grouped by gene_id. Final count matrices were annotated with gene biotype and metadata extracted from the reference GTF file. The 30 most differentially expressed genes were selected for downstream visualization and interpretation. MA plots were generated in RStudio using ggplot2, dplyr, and ggrepel packages.

*Flow Cytometry of Irradiated Organoids*

Prior to flow cytometry, organoids were dissociated to single cells using Neuron Dissociation Solutions (FujiFilm Biosciences, Cat# 297-78101) which included an Enzyme Solution (295-78021), an Dispersion Solution (298-78011) and an Isolation Solution (202-78031). Briefly, organoids were removed from culture and placed into 1.5mL tubes and were treated with each of the following solutions stepwise. Organoids were agitated at least 20 times in each solution and incubated at 37C for 5 minutes between agitation steps. After dissociation, supernatant was removed by centrifuging cells at 500g for 5 minutes at 4C. Cells were then washed with PBS and centrifuged again at 500g for 5 minutes at 4C. The resulting supernatant was removed, and the cells were incubated in PBS diluted with fixable viability dye eFluor450 (Invitrogen, Cat# 65-0863-15, 1:1000) for 15 minutes at room temperature to gate for live and dead cells. The cells were centrifuged at 500g for 5 minutes at 4C, and supernatant was removed. The cells were then permeablized and fixed using the BD Cytofix/Cytoperm™ Fixation/Permeabilization Kit (BD Biosciences, Cat#554714) according to the manufacturer's recommendations. After the removal of supernatant, cells were then washed once with FACS buffer solution. Cells were suspended in FACS buffer and run for flow cytometry using the Attune NxT Cytometer (Thermo Fisher Scientific). Data was analyzed using FCS Express (DeNovo Software).
